# Physicochemical and Antibacterial Evaluation of the methanol extract of *Chrysobalanus icaco* L. (Chrysobalanceae) Seed and Fractions

**DOI:** 10.64898/2025.12.12.680370

**Authors:** Emmanuel Oise Ikpefan, Jacinta Enoh Apitikori-Owumi, Lawrence Uchenna Nwankwo, David Enasawherie Ijamani

## Abstract

The use of plants as antimicrobial agents has gained popularity recently due to the need to explore safer alternative means of combating bacterial infections. The seeds of *C. icaco* were evaluated for its physicochemical, phytochemical and antimicrobial properties. The pulverized seed was extracted using cold maceration. Standard methods were used to evaluate the physicochemical properties of the powdered drug and phytochemical content of the crude extract of the seeds of *C. icaco*. The crude methanol extract was partitioned into dichloromethane and aqueous fractions. The antibacterial evaluation of the crude extract and fractions were done using agar well diffusion method. The phytochemicals present in the methanol crude were saponins, tannins, flavonoids, reducing sugars, alkaloids, steroid, terpenoids and cardiac glycosides. The moisture content value was 6.37 %. The antimicrobial result revealed that DCM fraction showed the highest zone of inhibition against *S. aureus, P. aeruginosa, S. typhi*, and *E. coli* at a concentration of 100 mg/mL, 200 mg/mL, 100 mg/mL, 25 mg/mL and 200 mg/mL respectively when compared to the crude seed extract and aqueous fraction. The MIC of crude extract was 50 mg/ml against each bacteria. The MIC of DCM fraction was between 100 mg/ml and 50 mg/ml against each bacteria. The MIC of aqueous fraction between 200 mg/ml and 50 mg/ml against each bacteria. This study therefore suggest that the crude extract and dichloromethane fraction of the seed of *C. icaco* possess antibacterial activities again selected isolates.

## 1. Introduction

Plants and plant components in the form of pressed, aqueous (cold, heated, or hot), and clandestine gin extracts have been used as dependable sources of medicine in traditional African civilizations and beyond, even before the arrival of orthodox medication.^1^. Public health is greatly concerned about the advent of bacteria that cause unknown diseases.^1^ This calls for updated approaches to both prevention and treatment, primarily through the creation of novel antibiotics.^2^ People are turning to homegrown medication due to the exorbitant expense of prescription drugs, the fragmentation of healthcare, and the availability of healing plants.^3^ Therefore, it is essential to do additional pharmacological research and development from any feasible source, and studying native flora may have promising outcomes. Numerous harmful diseases that affect people, animals, and plants are brought on by microorganisms. This is because bacteria become resistant to synthetic medications and antibiotics, they are unable to fully treat these diseases.^4^

*Chrysobalanus icaco* commonly known as “paradise plum”, “abajeru,” and “grageru.” “abajeru,” and “grageru” and is from the family of Chrysobalanaceae.^5^. It is classified as a medium-sized shrub.^6^ Locally it is called Omilo by the Iteskiri, Ebulo by the Ijaws and Amukan by the Yoruba.^5,7^ *Chrysobalanus icaco* is well known in Central America, Northwestern America, Carribean, South Florida, The Bahamas and in some West Africa countries such as Nigeria.^8,9^ Traditionally, *C. icaco* is used in the treatment of infections, leucorrhoea, diabetes, high cholesterol, inflammation, malaria, diarrhea and bleeding.^5,10^ According to Nojosa et al.^11^ *C. icaco* possesses biological activities that include anti-inflammatory, antibacterial, analgesic, anti-angiogenic, anti-diabetic, and anti-cancer effects.

One of the most significant and effective advances in contemporary science and technology for the prevention and treatment of infectious diseases is the discovery and development of antibiotics. Unfortunately, according to Habtamu *et al*.^12^, the pace at which pathogenic bacteria are becoming resistant to commonly used antimicrobial drugs is rising alarmingly. Globally, there is a rise in the isolation of microorganisms resistant to conventional antibiotics and the recovery of these isolates following antibacterial therapy.^12^ Moreover, antibiotics can have unfavorable side effects on the host, such as immunosuppression, hypersensitivity, loss of beneficial gut and mucosal microbes, and allergic reactions.^12^ In addition, the use of medicinal plants provides an excellent treatment option in developing countries all over the world. Hence scientific validations are essential for safe usage and improved effectiveness.^13^

Soups especially pepper soups are the primary means of *C. icaco* seeds consumption in Nigeria. The kernel that shakes freely within is taken out and pulverized. *Chrysobalanus icaco* fruits are edible, and they are frequently pickled sweetly in many different nations. The aim of the study is to evaluate the phytochemical profile and antimicrobial activities of the methanol extract and partitioned fractions of *Chrysobalanus icaco* seeds. The specific objective of this study includes extracting the seeds of *Chrysobalanus icaco* using a suitable solvent, investigating the phytochemical present in the methanol extract of *Chrysobalanus icaco* seeds, partitioning the methanol crude extract into aqueous and dichloromethane fractions, identifying the phytochemical constituents present and investigating the antimicrobial properties of the methanol extract and fractions of *Chrysobalanus icaco* seeds.

## 2. Methods

### 2.1 Reagents and Equipments

All solvents used were of Analar grade Methanol (JHD®, China), dichloromethane (DCM) (JHD®, China), distilled water, Dragendorff’s reagent, Mayer’s reagent, Fehling’s solution, Wagner’s reagent, Ferric chloride, NaOH, Olive oil, HCl, H_2_SO_4_, dilute ammonia solution, glacial acetic acid, dried seeds of *C. icaco*, cotton wool, beaker, test tubes, test tube racks, conical flask, measuring cylinder, Petri dishes, spatula, masking tape, glass rod, weighing balance (Shimadzu), ruler, refrigerator, sample bottle, separating funnel, rotary evaporator (Superfit Continental Ltd), desiccator, round bottom flask, laboratory incubator (Surgifriend Medicals, England), hot air oven (Leader Engineering St Helens Merseyside WAS), blender (FOOD MIXER, MODEL 300 C), porcelain dish, analytical balance (Shimadzu Model: ATY224 Philippines)), bijou bottles, electrical milling machine (Chris Morris, England), water bath (Labtop Instrument PVT, India), muffle furnace (Techmel & Techmel, Texas, USA), Whatman Grade 42 ashless filter paper, Whatman filter paper, (Whatmann International Ltd, Maidstone, England). Methanol (JHD®, China), dichloromethane (DCM) (JHD®, China), distilled water, Dragendorff’s reagent, Mayer’s reagent, Fehling’s solution, Wagner’s reagent, Ferric chloride, NaOH, Olive oil, HCl, H_2_SO_4_, dilute ammonia solution, glacial acetic acid.

### 2.2 Collection and identification of *Chrysobalanus icaco*

The fruits of the *C. icaco* were purchased from Warri Main Market Warri, Delta State, on December 16, 2023. The plant was authenticated at the Department of Plant Biology and Biotechnology, Herbarium Unit, Faculty of Life Sciences, University of Benin, Benin City, Edo State, Nigeria where the herbarium number UBH-C393 was obtained.

### 2.3 Preparation of the sample

The fruits of *C. icaco* were air-dried in the herbarium of the Department of Pharmacognosy and Traditional Medicine, Faculty of Pharmacy, Delta State University, Abraka for about 2 weeks to obtain a very dried fruit. The fruits were de-hulled to obtain the seeds which were dried, and pulverized into powder with a dried electronic blender.

#### 2.3.1 Seed extraction

About five hundred grams (500 g) of the resultant powdery specimen was extracted with about 2000 mL of methanol at room temperature (25±2 °C) for 3 days by cold maceration method with consecutive shaking. At the end of the 3^rd^ day, the extract was filtered through a Muslin cloth and was then concentrated to dryness on a rotary evaporator to form a semi-solid paste. The resulting crude extract was weighed with the weight recorded and then refrigerated till further use. The percentage yield of the methanol crude extracts of seeds of the *C. icaco* was calculated using the formula;

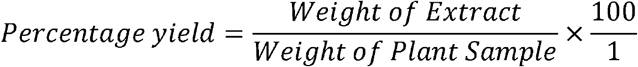

### 2.4 Fractionation of the methanol crude extract of the seed of *C. icaco*

The methanol seed extract was fractionated using a suitable grade of organic solvent (Dichloromethane and distilled water) in order to separate the active components present. The test was carried out using partitioning method in accordance with the method described by Akindele *et al*.^13^ with little modification. Thirty gram (30 g) of seed extract) was suspended in equal amount of methanol and distilled water (1:1), the mixture was then partitioned exhaustively with dichloromethane (DCM) (200 mL × 4) in a separating funnel (1L). The DCM fraction was collected successively at interval until partitioning was achieved. The aqueous and dichloromethane fractions obtained were concentrated to dryness and their weights were determined, labeled and stored in refrigerator at 4°C until required for use.

### 2.5 Qualitative phytochemical screening

Phytochemical Screening of the methanol crude extract and soluble partitioned fractions of *C. icaco* was done using the methods described by Dhivya.^14^ with little modifications.

### 2.6 Physicochemical evaluation of plant sample

The following criteria were used to determine the quality of the plant extract in accordance with recognized protocols.^15^ moisture content, ash value, extractive value, acid insoluble and water soluble ash was evaluated.

#### 2.6.1 Determination of moisture content

Two grams of the powdered sample were placed to the porcelain dish, and both the weight of the powdered sample and the porcelain dish were duly recorded after the porcelain dish was weighed. The powdered sample was placed in a porcelain plate and baked in a hot air oven. Using the temperature control knob, the hot air oven was brought up to 105 °C. The hot air oven was turned off after 15 minutes, and the porcelain dish with the dried extract was taken out and allowed to cool for another 15 minutes in a desiccator. After the dish holding the dried extract had cooled in the desiccator for fifteen minutes, it was weighed. Until a consistent weight is achieved, the drying, cooling, and weighing procedures are repeated. The moisture content of each plant extract is determined by calculating the difference between the final weight of the dried powdered plant and the porcelain dish and the initial weight of the powdered plant. The moisture content divided by the powdered plant’s starting weight and multiplied by 100 is the percentage moisture content.^15^

#### 2.6.2 Determination of total ash value

To eliminate all traces of moisture, an empty crucible was heated in a hot air furnace set to 105°C. After 20 minutes, the crucible was taken out of the hot air furnace and allowed to cool in a desiccator. The crucible’s weight was meticulously recorded. Each powdered sample was measured out to be 2 g in each crucible, which was then sealed with a lid. The temperature was adjusted to 700 °C and the two crucibles containing the powdered sample were placed inside the muffle furnace. The samples were left in the furnace for a duration of two hours. The muffle furnace was turned off after two hours, and the crucible was removed using a stainless steel locking tong and placed in a desiccator to cool. Following cooling, the crucible’s ultimate weight containing ash was recorded, and the percentage of ash was also computed.^15^

#### 2.6.3 Determination of acid insoluble ash

By mixing 40 mL of HCl with 60 mL of water, 40% HCl was created. After that, the 40% HCl was kept until needed in a glass bottle. After accurately measuring and adding 2 g of the powdered sample into the crucible, the crucible was set on a weighing balance. After being covered with a lid, the crucible was put inside a muffle furnace. For thirty minutes, the powdered sample was burned at 550°C. A few drops of distilled water were added to the ash content in the crucible to check for any potential development of black color, which usually indicates that the ash still has a lot of carbon. After 30 minutes, the crucible was removed from the furnace, the lid was removed, and the ash was carefully examined. The black color that developed after the distilled water was added made it necessary to burn once more. The crucible was set on a hot plate to properly dry the wet ash before it was put back in the furnace to burn.

When adding distilled water failed to produce any black color, the crucible containing the dry ash was put back into the furnace and left for an additional 40 minutes. At that point, it was discovered that the crucible was carbon free.

Following the verification of the ash’s carbon-free status, 25 milliliters of the 40% HCl solution were measured and transferred into the crucible that held the ash. By setting the crucible on the hot plate, the ash content in the HCl solution was brought to a boil for five minutes. Following boiling, the ash solution was filtered while still warm using an ashless filter paper setup.

The acid insoluble ash is defined as the residues that remain trapped on the filter paper during filtration. To guarantee that there is no remaining acid in the filter paper, the residue is removed from the filter using hot water.

After removing all traces of moisture, a crucible is cooled in a desiccator for ten minutes. It is first placed in a hot air oven for thirty minutes. After determining the crucible’s weight, the filter paper containing the acid-insoluble ash was put inside, the lid was put on, and the furnace was set for 90 minutes at 550 °C. After 90 minutes, the crucible was removed from the furnace and allowed to cool. At this stage, the acid insoluble ash in the crucible was weighed, and the percentage of acid insoluble ash was determined.^15^

#### 2.6.4 Determination of water soluble ash

A hot air oven was used to dry the crucible. The empty crucible and lid were weighed once they had cooled, and the weight was duly recorded. The crucible was filled with 2 g of the powdered sample, and the weight of the crucible, cover, and crude medication were also recorded. The crucible containing the crude drug was now put in the muffle furnace and allowed to get red hot. After that, it was turned off, and the crucible containing the ash was taken out and allowed to cool in the desiccator for twenty minutes. After weighing the crucible, lid, and ash, 25 mL of distilled water were added to dissolve the ash. The beaker was then filled with the ash-free solution and heated for fifteen minutes. Ashless filter paper was used to filter this; the filtrate comprises water soluble ash, while the filter paper holds on to the water insoluble ash. After being collected, the insoluble ash is put in a crucible and burned in an incinerator until it turns red hot. It is then carefully removed from the incinerator and put in a desiccator to cool. The difference between the water insoluble ash and the total ash is known as the water soluble ash. Finally, the appropriate percentage of water soluble ash was computed.^15^

#### 2.6.5 Determination of ethanol soluble extractives

Five grams of powdered drug were added to a glass container and then placed inside a dry 250 mL conical flask. A 100 mL graduated flask containing 90% ethanol was then poured into the conical flask containing the powdered drug. The conical flask was then corked, shaken constantly for the first six hours, and left to stand for the remaining eighteen hours. After twenty-four hours, when enough filtrate had been collected, filtering was done, and 25 mL of each filtrate were placed into a thin, weighed porcelain dish. On a water bath, the filtrate in the porcelain dish was evaporated until it was completely dry, and the drying process was finished in a hot air oven set at 100 °C. To guarantee cooling, the dried extract was thereafter placed in a desiccator. Weighing the cooled dried extract in the porcelain dish allowed us to determine the extractive’s weight percentage in relation to the air-dried medication.^15^

#### 2.6.5 Determination of water-soluble extractives

Five grams of powdered medication were placed to a glass container and then put into a dry 250 mL conical flask. A 100 mL graduated flask containing 99 mL of water and 1 mL of chloroform was filled with 99% distilled water. As a preservative, chloroform serves. The conical flask holding the powdered medication was filled with the mixture. After being corked and vigorously shaken for the first six hours, the conical flask is left to stand for the final eighteen hours. After a whole day, when enough filtrate has been gathered, filtration is completed. A thin porcelain plate was weighed and then filled with 25 mL of each filtrate. On a water bath, the filtrate in the porcelain dish was evaporated until it was completely dry, and the drying process was finished in a hot air oven set at 100 °C. To guarantee cooling, the dried extract was thereafter placed in a desiccator. Weighing the cooled dried extract in the porcelain dish allowed us to determine the extractive’s weight percentage in relation to the air-dried medication.^15^

#### 2.6.6 Determination of acetone, dichloromethane and methanol soluble extractives

Each powdered medication was weighed five grams in a weighing container before being put into a dry 250 mL conical flask. Three different substances were added to a 100 mL graduated flask: methanol, dichloromethane, and acetone. This was poured into the conical flask that held all of the powdered medications. After being corked and vigorously shaken for the first six hours, the conical flask is left to stand for the final eighteen hours. After 24 hours of filtering, 25 mL of each filtrate was placed to a thin, weighted porcelain plate once enough filtrate had been collected. On a water bath, the filtrate in the porcelain dish was evaporated until it was completely dry, and the drying process was finished in a hot air oven set at 100 °C. To guarantee cooling, the dried extract was thereafter placed in a desiccator. Weighing the cooled dried extract in the porcelain dish allowed us to determine the extractive’s weight percentage in relation to the air-dried medication.^15^

### 2.7 Isolation of bacteria

Isolation of bacteria was done using the method described by Hasan.^14^ and Biswas *et al*.^15^ with little modification. Organisms were isolated and enumerated by cultivating them on non-selective and selective media such as nutrient agar for total viable count (TVBC) and MacConkey agar for total coliform count (TCC). Mannitol salt agar was used for *Staphylococcus aureus*, Eosin methylene blue agar (EMB) for *Escherichia coli*, Centrimide agar for *Pseudomonas aeruginosa* and Samonella/Shigella agar for Samonella spp and Shigella spp. One loopful of culture media from nutrient broth was streaked over selective media and incubated overnight in an incubator at 37 ° C. Briefly, serial dilution method was use to isolate bacteria from the fresh fruits. Fresh fruits were pulverized in a sterile mortar and pestle along with distilled water to create a suspension, serially diluted up 10^1^ to 10^6^ times. A hundred microliter of diluted suspension of each of the fruit was spread over a well labelled solidified nutrient agar. The inoculated plates were incubated for 24 hours at 37 ° C.

#### 2.7.1 Macroscopic and microscopic identification of bacteria

The bacteria isolated were identified base on their biochemical and morphological characterization to Clauss ^16^ and Cheesbrough ^17^. Gram staining was carried out to study the cell morphology of the bacteria isolated. Every bacteria species were identified and confirmed via Indole production, Methyl-Red, Voges-Proskauer and Citrate (IMViC).^16,18^

### 2.8 Antibacterial activity

#### 2.8.1 Serial dilution of crude extract and fractions

A total of 15 test tubes were sterilized by autoclaving at 121 °C for 15 min and allowed to cool. For serial dilution of the crude extract, 2 mL of methanol was transferred into 4 test tubes, while 0.8 g of the crude extract was dissolved in 2 mL of methanol. Then 2 mL of the dissolved extract was transferred from the first test tube to the others using a 5 mL syringe and needle until the fifth test tube and 2 mL was discarded from the last test tube to give concentrations of 200 mg/mL, 100 mg/mL, 50 mg/mL, 25 mg/mL, 12.5 mg/mL. This same procedure was repeated for the dichloromethane (DCM) and aqueous fractions. The control used for bacteria was Ciprofloxacin (200 mg/100 mL).

#### 2.8.2 Antibacterial sensitivity testing of crude extract and fractions

The antibacterial activity of the plant extract was assessed using the agar cup diffusion method, as reported by Hugo ^19^. The Petri dishes were labelled according to the number of concentrations of extract, fractions and control. Sterilized molten Mueller Hinton agar was poured into Petri dishes for both the crude extract and fractions and rock gently for uniformity of thickness and allowed to solidify, the test was done in duplicates in aseptic conditions. The plates were inoculated with the different species of bacteria and hole was bored in the solidified media according to the concentrations and control. The various concentrations of extracts, fractions and control were transferred into the hole created. The plates were covered and left for 30 min to ensure diffusion of the crude extract and fractions. The plates inoculated with bacteria were incubated in the incubator at 37 ° C for 24 h. After 24 h of incubation, the plates were examined and the zones of inhibition were measured and recorded.

#### 2.8.3 Determination of the minimum inhibitory concentration of the crude extract and fractions

The plates were labeled according to the concentrations of the serial dilutions. Mueller Hinton agar was sterilized according to the manufacturer specification at 121 ° C for 15 min and allowed to cool. Each concentration was transferred into each plates for the various fractions and crude extract and the Mueller Hinton agar was poured into each plates containing the concentrations and gently rock for uniformity of thickness and allowed to solidify. The plates were inoculated with the various organisms that showed activity during the sensitivity test The plate streaked with bacteria were incubated at 37 ° C for 24 After 24 h, the minimum inhibitory concentration that inhibited the growth of organisms were determined.

### 2.9 Statistical analysis

Results were presented as mean ± standard error of the mean (SEM) of sample replicates. The data were analyzed using a one-way analysis of variance (ANOVA) followed by Dunnet’s test. *P*□0.05 were considered to be statistically significant

## 3. Results

### Percentage yield of crude extract and fractions of *C. icaco*

The percentage yield of the crude extract of *C. icaco* was 11.86% and the percentage yield of the partitioned fractions (dichloromethane and aqueous) of the crude extract of *C. icaco* were 43.33% and 11%, respectively. Table 1 showed the details of the percentage yield of the crude extract and partitioned fractions of *C. icaco*.

**Table 1:**
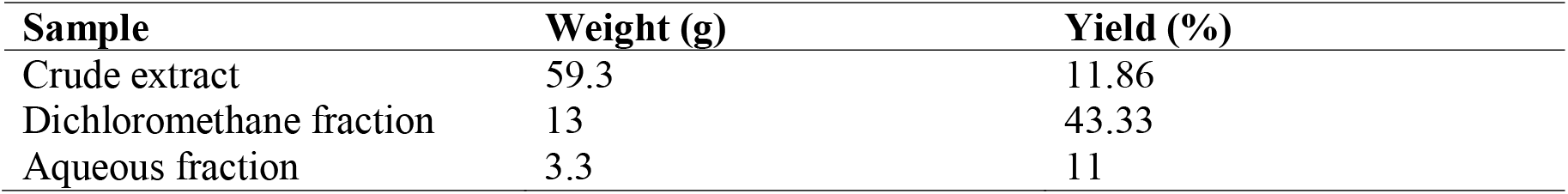
Percentage yield of crude extract and partitioned fractions of *C. icaco* seeds.

| Sample | Weight (g) | Yield (%) |
| --- | --- | --- |
| Crude extract | 59.3 | 11.86 |
| Dichloromethane fraction | 13 | 43.33 |
| Aqueous fraction | 3.3 | 11 |

### Physicochemical analysis

The results for physicochemical evaluation *C. icaco* seeds are presented in Table 2. Methanol, ethanol, and dichloromethane soluble extractive value (10.0%, 7.5% and 7.0%w/v, respectively) produced a higher yield of *Chrysobalanus icaco* seed extract when compared to acetone (6.5%) and water (5.0%) extractives.

**Table 2:**
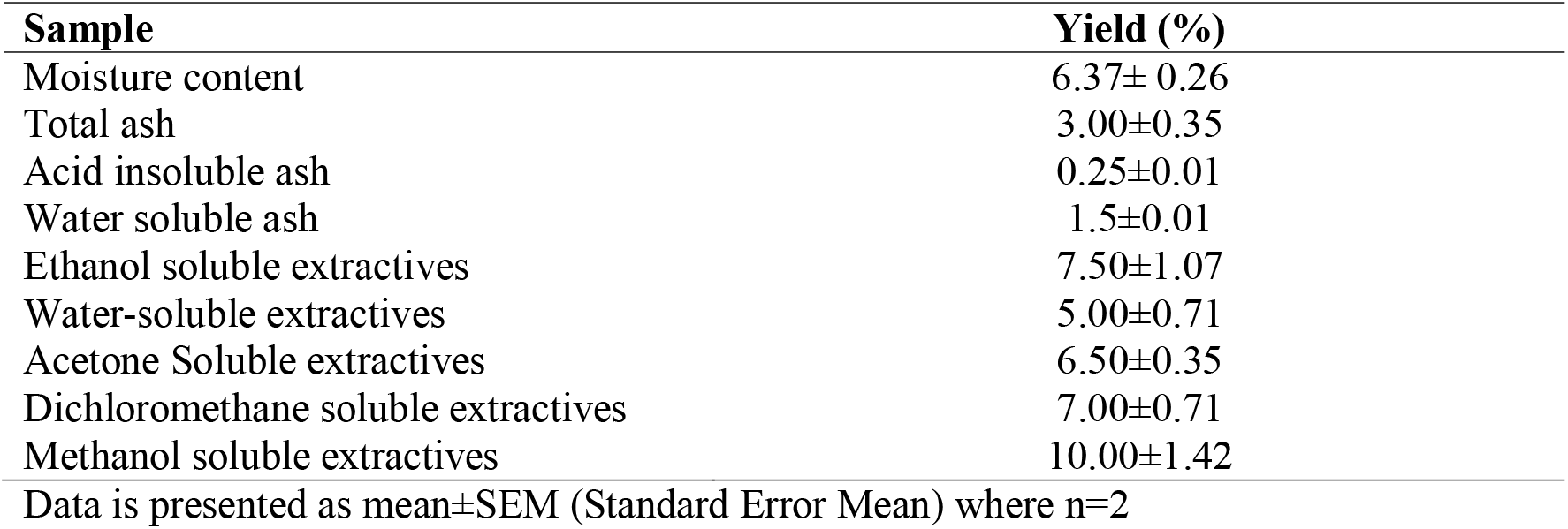
Result of the Physicochemical Evaluation of Powdered Sample of *C. icaco* seeds.

| Sample | Yield (%) |
| --- | --- |
| Moisture content | 6.37± 0.26 |
| Total ash | 3.00±0.35 |
| Acid insoluble ash | 0.25±0.01 |
| Water soluble ash | 1.5±0.01 |
| Ethanol soluble extractives | 7.50±1.07 |
| Water-soluble extractives | 5.00±0.71 |
| Acetone Soluble extractives | 6.50±0.35 |
| Dichloromethane soluble extractives | 7.00±0.71 |
| Methanol soluble extractives | 10.00±1.42 |

Data is presented as mean ± SEM (Standard Error Mean) where n=2

### Phytochemical analysis

The methanol seed extract of *C. icaco* were examined for the presence of secondary metabolites and the results are presented in Table 3 below. Eight metabolites were analyzed and the methanol crude extract showed the presence of seven metabolites which were saponin, tannin, flavonoid, reducing sugar, alkaloid, cardiac glycosides and terpenoid and the absence of one metabolite which was steroid.

**Table 3:** Results of the phytochemical screening of the leaf extract of *C. icaco* seeds.

| Phytochemical test | Methanol extract | Dichloromethane Fraction | Aqueous Fraction |
| --- | --- | --- | --- |
| Saponins | + | + | + |
| Tannins | + | + | + |
| Flavonoids | + | + | + |
| Steroids | - | + | - |
| Reducing sugars | + | - | + |
| Alkaloids | + | + | + |
| Terpenoids | + | + | + |
| Cardiac glycosides | + | + | - |

**Key:** + = Present, - = Absent

### Results for the antimicrobial activities of *C*.*icaco*

The antimicrobial activity of the methanol extract and partitioned fractions of *C*.*icaco* seeds against a panel of pathogenic microorganisms were evaluated. The results are presented in Tables 4 to 7. Table 4 showed antimicrobial activity of the methanol crude extract, dichloromethane and aqueous fractions of *C. icaco* seeds against *Staphylococcus aureus*. At 200 mg/mL, the dichloromethane fraction demonstrated higher inhibition (10.00±0.71 mm) than crude methanol extract (7.50±1.78 mm), and aqueous fraction (4.00±0.00 mm). At 100 mg/mL, the dichloromethane fraction also demonstrated higher inhibition (15.5±0.35) than crude methanol extract (4.00±0.71mm), and the aqueous fraction (2.00±0.00 mm). Moreover, at 50 mg/mL, the dichloromethane fraction also demonstrated higher inhibition (2.00±1.42 mm) than crude methanol extract (1.00±1.42 mm), while the aqueous had no inhibition. In addition, at 25 mg/mL, the crude methanol extract demonstrated inhibition (0.50±0.35 mm), while the dichloromethane and aqueous fractions had no inhibition.

**Table 4:**
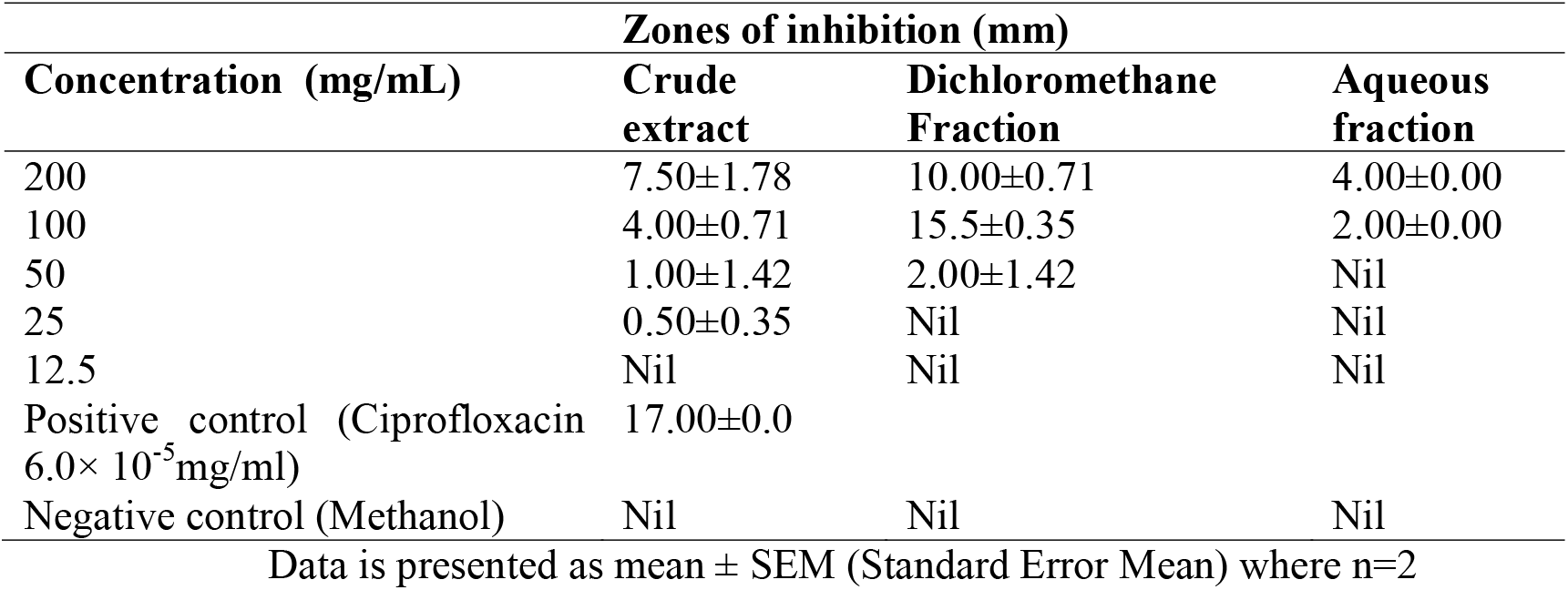
Antimicrobial activity of the methanol crude extract, dichloromethane and aqueous fractions of *C. icaco* seeds against *Staphylococcus aureus*.

| Concentration (mg/mL) | Zones of inhibition (mm) |  |  |
| --- | --- | --- | --- |
|  | Crude extract | Dichloromethane Fraction | Aqueous fraction |
| 200 | $7.50 \pm 1.78$ | $10.00 \pm 0.71$ | $4.00 \pm 0.00$ |
| 100 | $4.00 \pm 0.71$ | $15.5 \pm 0.35$ | $2.00 \pm 0.00$ |
| 50 | $1.00 \pm 1.42$ | $2.00 \pm 1.42$ | Nil |
| 25 | $0.50 \pm 0.35$ | Nil | Nil |
| 12.5 | Nil | Nil | Nil |
| Positive control (Ciprofloxacin $6.0 \times 10^{-5}$ mg/ml) | $17.00 \pm 0.0$ | | |
| Negative control (Methanol) | Nil | Nil | Nil |

Data is presented as mean ± SEM (Standard Error Mean) where n=2

Table 5 showed that *Pseudomonas aeruginosa* was susceptible to the crude extract, dichloromethane fraction and slightly susceptible to aqueous fraction *of C. icaco*. At 200 mg/mL, the dichloromethane fraction demonstrated higher inhibition (16.00±4.28 mm) than crude methanol extract (10.00±2.85 mm), while the aqueous fraction had the least inhibition (2.00±0.71mm). At 100 mg/mL, the dichloromethane fraction also demonstrated higher inhibition (8.00±0.71) than crude methanol extract (7.50±1.78mm), while the aqueous fraction exhibited zero inhibition. Moreover, at 50 mg/mL, the dichloromethane fraction also demonstrated higher inhibition (14.00±2.85 mm) than crude methanol extract (2.00±1.42 mm), while the aqueous had no inhibition. In addition, at 25 mg/mL, the dichloromethane fraction demonstrated higher inhibition (9.50±1.78 mm), while the crude methanol extract and aqueous fractions had no inhibition. At 25 mg/mL, the dichloromethane fraction demonstrated inhibition (9.50±1.78 mm), while the crude methanol extract and aqueous fractions had no inhibition.

**Table 5:** Antimicrobial activity of the methanol crude extract, dichloromethane and aqueous fractions of *C. icaco* seeds against *Pseudomonas aeruginosa*.

| Concentration (mg/mL) | Zones of inhibition (mm) |  |  |
| --- | --- | --- | --- |
|  | Crude extract | Dichloromethane fraction | Aqueous fraction |
| 200 | $10.00 \pm 2.85$ | $16.00 \pm 4.28$ | $2.00 \pm 0.71$ |
| 100 | $7.50 \pm 1.78$ | $8.00 \pm 0.71$ | Nil |
| 50 | $2.00 \pm 1.42$ | $14.00 \pm 2.85$ | Nil |
| 25 | Nil | $9.50 \pm 1.78$ | Nil |
| 12.5 | Nil | $15 \pm 0.71$ | Nil |
| Positive control (Ciprofloxacin) | $20.00 \pm 2.85$ | $7.00 \pm 5.00$ | Nil |
| Negative control | Nil | Nil | Nil |

Data is presented as mean ± SEM (Standard Error Mean) where n=2

The results obtained in Table 6 showed that *Salmonella typhi* was susceptible to the crude extract, dichloromethane fraction and the aqueous fraction *of C*.*icaco*. At 200 mg/mL, the aqueous fraction demonstrated higher inhibition (9.50±1.78 mm) than dichloromethane fraction (9.00±0.71mm), while the crude methanol extract had the least inhibition (5.00±0.71mm). At 100 mg/mL, the dichloromethane fraction also demonstrated higher inhibition (10.50±1.78 mm) than crude methanol extract (5.50±0.125 mm), while the aqueous fraction had the least inhibition (2.50±0.35 mm). Moreover, at 50 mg/mL, the dichloromethane fraction also demonstrated higher inhibition (6.00±0.00 mm) than crude methanol extract (3.00±0.71 mm), while the aqueous fraction demonstrated the least inhibition (2.50±0.35 mm). In addition, at 25 mg/mL, the dichloromethane fraction demonstrated higher inhibition (9.50±1.78 mm), while the crude methanol extract and aqueous fractions had no inhibition. At 25 mg/mL, the dichloromethane fraction demonstrated a higher inhibition (4.50±0.71 mm) compared to the aqueous fraction (2.50±0.35), while the crude methanol extract had no inhibition. At 12.5 mg/mL, the dichloromethane fraction demonstrated a higher inhibition (3.50±0.35 mm) compared to the aqueous fraction (1.50±0.35 mm), while the crude methanol extract had no inhibition.

**Table 6:**
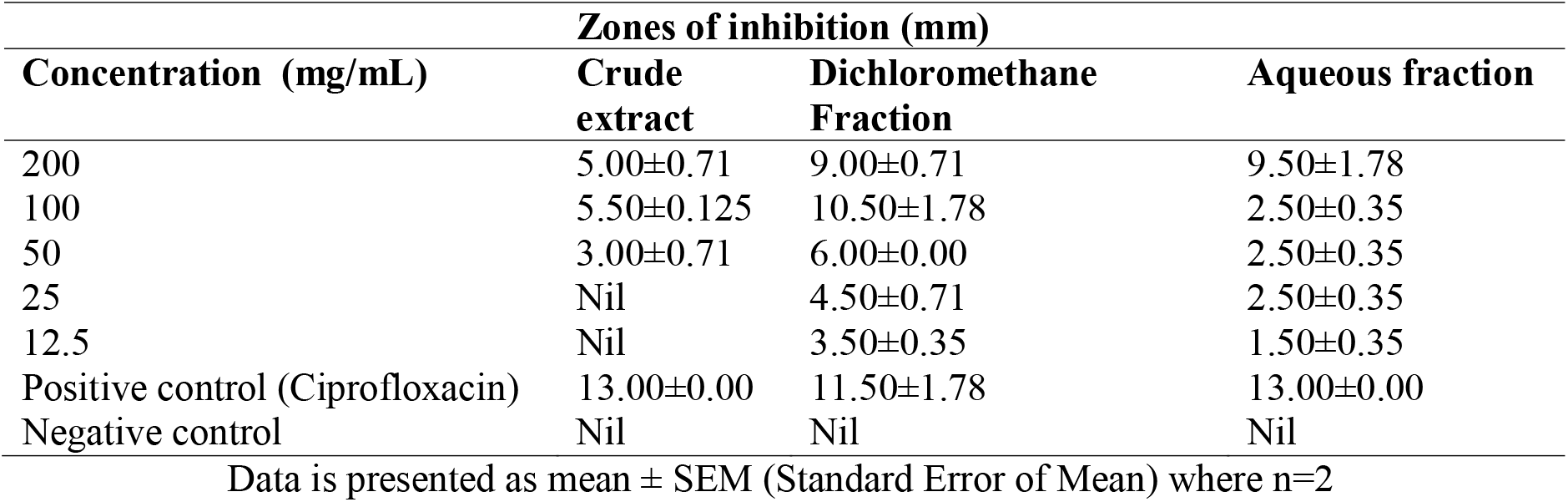
Antimicrobial activity of the methanol crude extract, dichloromethane and aqueous fractions of *C. icaco* seeds against *Salmonella typhi*.

The results obtained in Table 7 show that *Escherichia coli* was susceptible to the crude extract, dichloromethane fraction and the aqueous fraction *of C*.*icaco*. At 200 mg/mL, the dichloromethane fraction demonstrated higher inhibition (8.5±1.07 mm) than crude methanol extract (4.5±0.35 mm), while the aqueous fraction had the least inhibition (2.5±0.35 mm). At 100 mg/mL, the dichloromethane fraction also demonstrated higher inhibition (11.5±0.35 mm) than crude methanol extract (2.5±1.07 mm), while the aqueous fraction had the least inhibition (2±0.00 mm). Moreover, at 50 mg/mL, the dichloromethane fraction also demonstrated higher inhibition (11±0.71 mm) than aqueous fraction (3.5±0.35 mm), while the crude methanol extract demonstrated no inhibition. Moreover, at 25 mg/mL, the dichloromethane fraction demonstrated higher inhibition (13.5±0.35 mm) than aqueous fraction (2±0.00 mm), while the crude methanol extract demonstrated no inhibition. At 12.5 mg/mL, the dichloromethane fraction also showed an inhibition (10±0.71 mm), while the crude methanol extract and aqueous fraction had no inhibition.

**Table 7.** Antimicrobial activity of the methanol crude extract, dichloromethane and aqueous fractions of *C. icaco* seeds against *Escherichia coli*.

| Concentration (mg/mL) | Zones of inhibition (mm) |  |  |
| --- | --- | --- | --- |
|  | Crude extract | Dichloromethane Fraction | Aqueous fraction |
| 200 | 4.5±0.35 | 8.5±1.07 | 2.5±0.35 |
| 100 | 2.5±1.07 | 11.5±0.35 | 2±0.00 |
| 50 | Nil | 11±0.71 | 3.5±0.35 |
| 25 | Nil | 13.5±0.35 | 2±0.00 |
| 12.5 | Nil | 10±0.71 | Nil |
| Positive control (Ciprofloxacin) | 15.50±0.35 | 14.5±0.35 | 14.5±0.35 |
| Negative control | Nil | Nil | Nil |

Data is presented as mean ± SEM (Standard Error Mean) where n=2

### Results of minimum inhibitory concentration

The minimum inhibitory concentration of the methanolic extract and partitioned fractions of *C. icaco* seeds against a panel of pathogenic microorganisms were evaluated. The results are presented in Tables 8 to 10.

**Table 8:** Minimum inhibitory concentration of the methanol crude extract of *C. icaco* seeds against the different test organisms.

| Organisms | Concentration (mg/mL) |  |  |  |  |
| --- | --- | --- | --- | --- | --- |
|  | 200 | 100 | 50 | 25 | 12.5 |
| *Staphylococcus aureus* | - | - | - | + | + |
| *Escherichia coli* | - | - | - | + | + |
| *Salmonella typhi* | - | - | - | + | + |
| *Pseudomonas aeruginosa* | - | - | - | + | + |

**Table 9:** Minimum inhibitory concentration of the dichloromethane fraction of *C. icaco* seeds against the different test organisms.

| Organisms | Concentration (mg/mL) |  |  |  |  |
| --- | --- | --- | --- | --- | --- |
|  | 200 | 100 | 50 | 25 | 12.5 |
| *Staphylococcus aureus* | - | - | + | + | + |
| *Escherichia coli* | - | - | + | + | + |
| *Salmonella tspp* | - | - | - | + | + |
| *Pseudomonas aeruginosa* | - | - | - | + | + |

Key: -= No growth, += Growth

**Table 10:** Minimum inhibitory concentration of the aqueous fraction of *C. icaco* seeds against the different test organisms.

| Organisms | Concentration (mg/mL) |  |  |  |  |
| --- | --- | --- | --- | --- | --- |
|  | 200 | 100 | 50 | 25 | 12.5 |
| *Staphylococcus aureus* | - | - | - | + | + |
| *Escherichia coli* | - | + | + | + | + |
| *Salmonella typhi* | - | + | + | + | + |
| *Pseudomonas aeruginosa* | - | + | + | + | + |

### Results of minimum inhibitory concentration

The minimum inhibitory concentration of the methanol extract and partitioned fractions of *C*.*icaco* seeds against a panel of pathogenic microorganisms were evaluated. The results are presented in Tables 11 to 13.

**Table 11:**
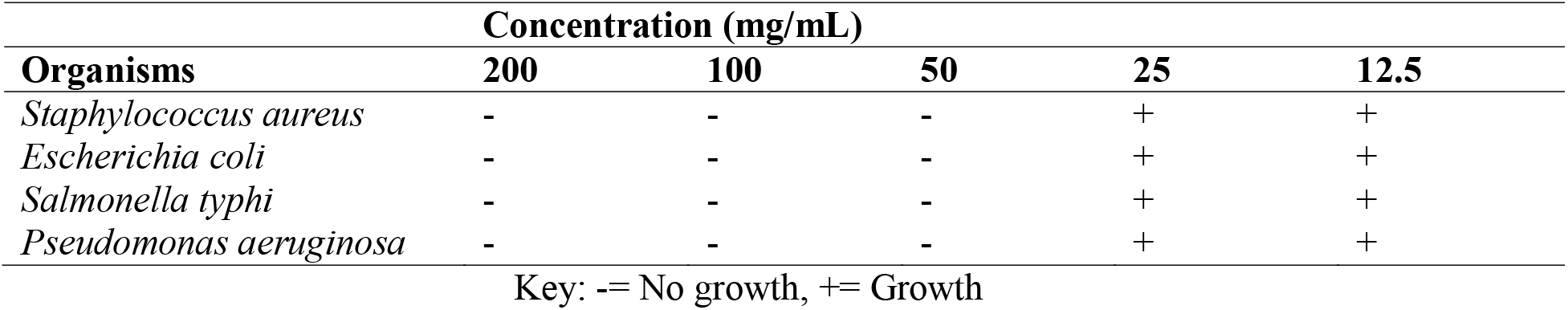
Minimum inhibitory concentration of the methanol crude extract of *C. icaco* seeds against the different test organisms.

**Table 12:** Minimum inhibitory concentration of the dichloromethane fraction of *C. icaco* seeds against the different test organisms.

| Organisms | Concentration (mg/mL) |  |  |  |  |
| --- | --- | --- | --- | --- | --- |
|  | 200 | 100 | 50 | 25 | 12.5 |
| *Staphylococcus aureus* | - | - | + | + | + |
| *Escherichia coli* | - | - | + | + | + |
| *Salmonella typhi* | - | - | - | + | + |
| *Pseudomonas aeruginosa* | - | - | - | + | + |

**Table 13:**
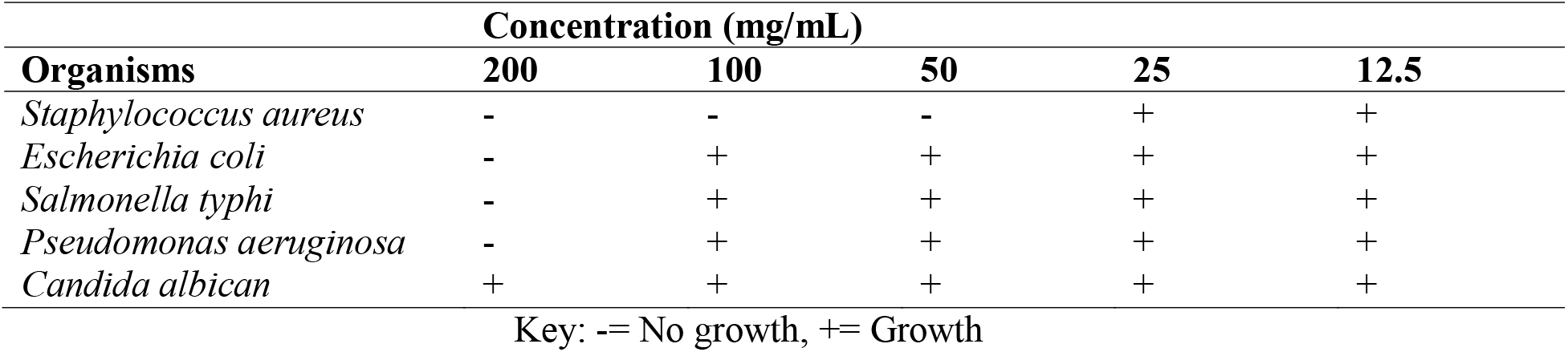
Minimum inhibitory concentration of the aqueous fraction of *C. icaco* seeds against the different test organisms.

| Organisms | Concentration (mg/mL) |  |  |  |  |
| --- | --- | --- | --- | --- | --- |
|  | 200 | 100 | 50 | 25 | 12.5 |
| *Staphylococcus aureus* | - | - | - | + | + |
| *Escherichia coli* | - | + | + | + | + |
| *Salmonella typhi* | - | + | + | + | + |
| *Pseudomonas aeruginosa* | - | + | + | + | + |
| *Candida albican* | + | + | + | + | + |

Key: -= No growth, += Growth

## 4. Discussion

Plants are one of the cornerstones for modern medicine to achieve new principles because of the potential for producing antimicrobials from higher plants, which would result in the development of a medication to work against bacteria. The percentage yield of the *Chrysobalanus icaco* methanol seed extract was quite minimal (11.86%). One of the following may be responsible, the surface area, the moisture content of the crude drug, the poor diffusion rate of the powdered drug, or the existence of competing extractable components. The type of the extraction solvent or an unbalanced liquid-solid ratio might also be blamed.

The physical constants employed in the assessment of crude pharmaceuticals are crucial metrics for identifying instances of medication adulteration or inappropriate storage (DanMallam *et al*., 2001). The moisture content of *Chrysobalanus icaco* seeds obtained in this study was 6.37± 0.26 which is lower than the standard requirement of moisture in crude drugs (8-14%) according to the African Pharmacopooeia ^20^, this indicates that the moisture content is low and there may be very minimal or no tendency of microbial degradation or spoilage. The moisture content is very significant in terms of prolonged shelf life and resistance to microbial degradation of the crude drug ^21^.

For the assessment of a crude drugs, extractive values are helpful. It provides insight on the kind of chemical components found in the crude drugs. It is helpful for estimating the components extracted using the extraction solvent ^22^. Drugs that are depleted or tampered with can be identified using extractive values. Both the drug’s grade and purity are determined by the extractive value of the crude ^22^. The extractive values of the crude drug using water, ethanol, methanol, dichloromethane, and acetone were thus calculated. To accurately estimate which solvent is most likely to extract a higher yield of phytoconstituents, a variety of solvents were utilized. According to the study by Imieje and Ezenwanne ^21^, it was discovered that the extractive values of alcohol (methanol, ethanol) were higher than those of water. This suggests that alcohol may be a more effective solvent for extraction than water.

When identifying foreign inorganic materials that are present as contaminants in crude medications, ash values can be helpful. The *C. icaco* seeds had an overall ash content of 3.00±0.35, indicating a low degree of contaminants in the crude medication. This result is comparable to the total ash content value that Silva *et al*. ^8^ previously ascertained. According to Chaudhari and Girase ^22^, the drug’s purity, inorganic composition, and presence of foreign materials were indicated by the values of water soluble ash and acid insoluble ash. *C. icaco*’s low levels of water soluble ash (1.5±0.01) and acid insoluble ash (0.25±0.01) indicated that the sample was devoid of any foreign material. This is consistent with the findings of Silva *et al*. ^8^ Phytochemicals present in the crude methanol extract and the fractions of *C. icaco* seeds were alkaloids, saponin, cardiac glycosides, terpenoids, tannins, flavonoids, and phenol while steroids were not present. Phytochemicals have been known to contribute immensely to the antibacterial properties of most crude drugs. Recent studies have shown that they are vital in addressing issues of bacterial resistance.^23^

The result of the study revealed that the methanol crude extract of *C. icaco* seed, as well as the dichloromethane and aqueous fractions were all effective against *S. aureus*.

Similarly, the results obtained in this study also showed that *S. typhi, E. coli* and *Pseudomonas aeruginosa* were susceptible to the crude extract, dichloromethane fraction and slightly susceptible to aqueous fraction *of C. icaco*. Terpenoids, alkaloids and saponins inhibit bacteriostatic and bactericidal activity by altering the physiological make-up of the bacteria. This anomaly leads to the loss of important proteins and enzymes, thus achieving antimicrobial effect.^24^ The findings of this study and previous study clearly points out the presence of these aforementioned phytochemicals, hence they can be attributed to the susceptibility of the microorganisms towards the crude extract and the dichloromethane fractions.

The antibacterial compounds in many crude drugs are often non-polar or moderately polar, making them more soluble in organic solvents like dichloromethane than water. The aqueous fractions obtained from this study may have isolated more of phytochemicals such as tannins and proteins which may not possess strong antibacterial activity. Some antibacterial compounds may be unstable or susceptible to degradation when exposed to water, this could be as a result of hydrolysis or oxidation. The crude extract’s antibacterial activity in comparison to the aqueous extract is due to the fact that it contains a wide range of compounds that works synergistically to enhance antibacterial effects. Ciprofloxacin is known to exhibit a high level of efficacy against wide range of gram positive and gram negative bacteria via inhibition of DNA gyrase enzyme responsible for DNA supercooling and subsequent bacteria DNA replication. In this study, the positive control drug (Ciprofloxacin) exhibited significantly higher antibacterial activity against test isolates when compared with the crude methanol extract and fractionated extracts.

In this study, the MIC of the methanol crude extract from *C. icaco* seeds against *S*. aureus, *Escherichia coli, S. typhi*, and *P. aeruginosa* was 50 mg/mL. However, the MIC of the dichloromethane partitioned fraction from the same seeds was 100 mg/mL for *S. aureus* and *E. coli*, and 50 mg/mL for *S. typhi* and *P. aeruginosa*. The results of the minimum inhibitory concentration (MIC) of the aqueous partitioned fraction of *C. icaco* seeds against *S. aureus* was 50 mg/mL, while that of *Escherichia coli, S. typhi and P. aeruginosa* was 200 mg/mL. These results contradicts the findings of Castilho and Kaplan ^25^ where the MIC for *S. aureus* and *S. pyogenes* was about 60 μg/mL.

## Conclusion

This study therefore suggest that the crude methanol extract of the seed of *C. icaco* and its dichloromethane fraction possess good antibacterial effect against selected isolates. The aqueous fractions on its part, demonstrated feeble effect against same isolates thereby expressing its limitation as a suitable antibacterial agent against the tested isolates.

## Acknowledgements

All authors express their gratitude to God for the success of this work.

## Authors’ Contributions

IEO and AJE conceptualized the initial idea and designed the scope and directions of the study. AJE, NLU and IDE made equal contributions to different part of the review. NLU contributed to the review and managed the references. AJE and IDE designed, retrieved and analyzed the survey and data obtained. AJE and NLU thoroughly overhauled the manuscript and made valuable inputs. All authors read and edited the final copy of the manuscript. IEO gave final authorization for the submission of the manuscript.

## References

1. Onilude HA, Kazeem MI and Adu OB (2021) Chrysobalanus icaco: A review of its phytochemistry and pharmacology. Journal of Integrative Medicine 19(1): 13–19

2. Iwu MM (2019) Handbook of African Medicinal Plants. CRC.Press Boca Baton, pp.11–15

3. Rasyid ZA, Farida A, Daud SH, Wiwin, S, Wijaya, KI and Tangke, AE (2020) Bioactivities of Forest Medicinal Plants on Kutai Ethnic (Indonesia) of Tapak Leman (Hippobroma longiflora (L) G. Don). GSC Biological and Pharmaceutical Sciences 11:91–98

4. Nojosa ECN, Marques GEC, Dos Anjos da Paz S, Reis, JT, Brandão, CM, do Nascimento, AS, Camara, MBP and Dos Santos DR (2023) Bioactivity and antimicrobial evaluation of extracts from Chrysobalanus icaco L. found in the Amazonian maranhense, Brazil. Revista Gestão e Secretariado (GeSec). Journal of Management and Secretarial Science 14 (9);15537–15551.

5. Apitikori-Owumi JE, Sonibare MA, Marwe AA and Firdous SM (2024) An Ethnobotanical Survey, Pharmacognostic Profile and Phytochemical Analysis Investigation of Chrysobalanus icaco L. Yuzuncu Yil University Journal of Agricultural Sciences 34(3): 489–504.

6. Kadiri HE (2019) The modulatory effect of Chrysobalanus icaco seed on cadmium induced hepatoxicity and nephrotoxicity in rats. Nigerian. Journal of Pure and Applied Science 32(2):3428–3435.

7. Erhenhi AH, Lemmy EE and Ashibuogwu CC (2016) Spices used in Ubulu-Uku community of Delta state. International Journal of Herbal Medicine 4(3):45

8. Silva JP, Peres AM, Paixão TP, Silva AS, Baetas AC and Barbosa WL (2017) Antifungal activity of hydroalcoholic extract of Chrysobalanus icaco against oral clinical isolates of Candida Species. Pharmacognosy Research 9: 96–100.

9. Adomi PO, Nana MJ and Owhe-Ureghe UB (2024) Phytochemical properties and antibacterial effects of Aframomum sceptrum, Chryosobalanus icaco and Piper guineense seeds. World Journal Of Pharmaceutical and Medical Research 10(7): 45–49

10. Araújo-Filho HG, Dias JDS, Quintans-Júnior LJ, Santos MRV, White PAS and Barreto RSS (2016) Phytochemical screening and analgesic profile of the lyophilized aqueous extract obtained from Chrysobalanus icaco leaves in experimental protocols. Pharmaceutical Biology 54:3055–3062.

11. Nojosa, ECN, Marques, GEC, dos Anjos da Paz, S, Reis, JT, Brandão, CM, do Nascimento, AS, Camara, MBP and dos Santos, D. (2023) Bioactivity and antimicrobial evaluation of extracts from Chrysobalanus icaco L. found in the Amazonian maranhense, Brazil. Revista Gestão e Secretariado 14(9): 15537–15551.

12. Habtamu T, Velmurugan S, Addisu A (2018) A Review on Antimicrobial Activity of Medicinal Plants against Human Pathogens. Journal of Natural Science Research 8(19):10–19

13. Akindele EO, Ehlers SM and Koop JHE (2019) First empirical study of freshwater microplastics in West Africa using gastropods from Nigeria as bioindicators. Limnologica 78: 9

14. Dhivya R and Manimegalai K (2013) Preliminary Phytochemical Screening and GC-MS Profiling of Ethanolic Flower Extract of Calotropis gigantea Linn. (Apocynaceae). Journal of Pharmacognosy and Phytochemistry 2(3):28–32.

15. Nwankwo LU, Obokare EC (2023). Preliminary Phytochemical, Physicochemical, and Comparative Antibacterial Evaluation of Methanolic Extracts of Ocimum gratissimum Stem against Methicillin-Resistant Staphylococcus aureus and Proteus vulgaris. International Pharmacy Acta 2023: 6(1):e8

16. Clauss D and Berkeley RCW (1986) Genus Bacillus In Bergey’s manual of determinative bacteriology, Sneath P H A Baltimore. MD: Williams Wilkins, 2, 1105

17. Cheesbrough M (2004). District Laboratory Practice in Tropical Countries part 2. Low price editions. Cambridge University press, London. p 434

18. Hasan NA and Zulkahar IM (2018) Isolation and identification of bacteria from spoiled fruits. AIP Conference Proceedings 1: 020073

19. Hugo WB and Russel AD (1992) Pharmaceutical Microbiology, 5th Edition, Blank Well Scientific Publication, Oxford, London. pp. 258–297

20. African Pharmacopoeia (1986) General Methods for Analysis. OAU/SRTC Scientific Publications Lagos, pp. 137–149.

21. Rezaei F and VanderGheynst JS (2010) Critical Moisture content for microbial growth in dried food-processing residues. Journal of the science of food and Agriculture 90(12): 2000–2005. Doi:10.1002/jsfa.4044

22. Chaudhari RK and Girase NO (2015) Determination of soluble extractives and physico-chemical studies of bark of Sesbania sesban (L) Merr. Journal of Chemical and Pharmaceutical Research 7(8):657–660.

23. Khameneh B, Eskin NAM, Iranshahy M and Bazzaz BSF (2021) Phytochemicals: A Promising weapon in the Arsenal against Antibiotic –Resistant Bacteria. Antibiotics (Basel), 10(9):1044. Doi:10.3390/antibiotics10091044

24. Huang W, Wang Y, Tian W, Cui X, Tu P, Li J, Shi S and Liu X (2022) Biosynthesis Investigations of Terpenoid, Alkaloid, and Flavonoid Antimicrobial Agents Derived from medicinal plants. Antibiotics (Basel) 11(10):1380. Doi:10.3390/antibiotics11101380

25. Castilho RO and Kaplan MAC (2011) Phytochemical study and antimicrobial activity of Chrysobalanus icaco. Chemistry of Natural Compounds 47:436–437.

